# Atlas of the Bacterial Serine-Threonine Kinases expands the functional diversity of the kinome

**DOI:** 10.1101/2025.01.12.632604

**Authors:** Brady O’Boyle, Wayland Yeung, Jason D. Lu, Samiksha Katiyar, Tomer M. Yaron-Barir, Jared L. Johnson, Lewis C. Cantley, Natarajan Kannan

## Abstract

Bacterial serine-threonine protein kinases (STKs) regulate diverse cellular processes associated with cell growth, virulence, and pathogenicity. They are evolutionarily related to the druggable eukaryotic STKs. However, an incomplete knowledge of how bacterial STKs differ from their eukaryotic counterparts and how they have diverged to regulate diverse bacterial signaling functions presents a bottleneck in targeting them for drug discovery efforts. Here, we classified over 300,000 bacterial STK sequences from the NCBI RefSeq non-redundant and UniProt protein databases into 35 canonical and seven non-canonical (pseudokinase) families based on the patterns of evolutionary constraints in the conserved catalytic domain and flanking regulatory domains. Through statistical comparisons, we identified distinguishing features of bacterial STKs, including a distinctive arginine residue in a regulatory helix (C-Helix) that dynamically couples ATP and substrate binding lobes of the kinase domain. Biochemical and peptide-library screens demonstrated that constrained residues contribute to substrate specificity and kinase activation in the *Mycobacterium tuberculosis* kinase PknB. Collectively, these findings open new avenues for investigating bacterial STK functions in cellular signaling and for the development of selective bacterial STK inhibitors.

## Introduction

Hanks-type serine-threonine protein kinases, often referred to simply as STKs or protein kinases, are a large family of enzymes that play crucial roles in metabolism, cell signaling, protein regulation, cellular transport, secretory processes, and other cellular pathways (*1–3*). They accomplish this by catalyzing the transfer of a phosphate group - usually from ATP - to a serine or threonine residue in a target protein substrate, thereby regulating protein functions post translationally. Phosphorylation by serine-threonine protein kinases is a prevalent form of regulation found across all domains of life (*4–8*). The human genome encodes ∼450 STKs (*1*), and orthologs of these proteins have been mapped across diverse eukaryotic species (*2*, *9*). Further evolutionary classification of these sequences based on the patterns of conservation and variation in the commonly conserved catalytic domain has identified sequence and structural features contributing to the functional specialization of STK families (*10–18*) as well as in the development of selective STK inhibitors (*19–23*).

Despite substantial progress in the evolutionary and functional characterization of eukaryotic serine-threonine protein Kinases (eSTKs), the STK complement of microbial genomes remains relatively understudied. One group of understudied Hanks-type protein kinases is the bacterial serine-threonine protein Kinase (bSTK) class, sometimes collectively referred to as PknB (*24*, *25*). bSTKs are unrelated to bacterial tyrosine kinases or histidine kinases (*26*, *27*), but share a high degree of sequence and structural similarity to eSTKs (*28*). Initially discovered in *Myxococcus xanthus* in 1991 (*29*), the number of STKs within bacterial genomes ranges from 1 in *Escherichia coli* to over 60 in some species of Actinobacteria (*25*). bSTKs are of particular interest because of their role in virulence (*30–40*), bacterial cellular development (*30–32*, *35*, *41–50*), and antibiotic resistance (*30*, *51–56*). A greater understanding of bSTKs and their integral role in bacterial cellular processes can shed light on their potential use as targets for antibacterial drugs (*57–61*), further evidenced in a study where a bSTK was targeted to successfully inhibit the growth of methicillin-resistant *Staphylococcus aureus* (MRSA), a bacterial strain that is resistant to many commonly used antibiotics (*56*).

In particular, the extensive library of small molecule inhibitors developed for eSTKs can potentially be repurposed for targeting bSTKs involved in bacterial virulence and antibiotic resistance. However, an incomplete knowledge of the evolutionary divergence of bSTKs and eSTKs, and the lack of an evolutionary framework relating diverse bSTK families, has hindered drug discovery efforts and structural understanding of bSTK functional specialization. Previous efforts to identify and classify bSTK sequences have primarily focused on a subset of prokaryotic phyla (*25*, *62*, *63*). Although these studies have illuminated pieces of the diversity and evolution of bSTKs, there has yet to be any unified, systematic classification or evolutionary analysis of bSTKs to the degree seen with human protein kinases.

Here, we classified nearly 300,000 bSTK protein sequences spanning nearly 26,000 strains into 42 distinct families comprising canonical kinase and noncanonical pseudokinase families, homologous to Pfam pkinase family (PF00069) (*64*). Furthermore, using a Bayesian statistical approach successfully employed in the functional classification of eSTKs (*11*, *16–18*, *65–71*), we delineated sequence constraints characteristic of various bSTK families. We analyzed them using available crystal structures and AlphaFold models. In addition to unique domain combinations and sequence indels, we identified co-conserved sequence patterns characteristic of various bSTK families. Biochemical and peptide-library-based characterization of a bSTK conserved RF motif in the C-Helix of the *Mycobacterium tuberculosis* kinase PknB revealed new insights into substrate recognition and allosteric control of catalytic activity. By revealing shared and unique regulatory mechanisms for bSTKs, our studies open new avenues for repurposing eSTK drugs for bacterial infections.

## Results

### 42 bSTK families identified through constraint-based clustering

We identified bSTK families using constraint-based sequence clustering of aligned protein sequences from the UniProt (https://www.uniprot.org) and NCBI RefSeq non-redundant (https://www.ncbi.nlm.nih.gov/refseq/about/nonredundantproteins/) protein sequence databases. To ensure an accurate sequence alignment of bSTK catalytic domains, we began by generating an alignment profile using 43 experimentally validated bSTKs (data file S1). We then utilized multiply-aligned profiles for global alignment of protein sequences (MAPGAPS) (*72*), a profile-based alignment tool, to identify, align and annotate bSTKs from the wealth of publicly available sequence data in the RefSeq Non-Redundant protein sequence database (NR) (*73*) and UniProtKB + UniRef protein sequence database (*74*). We then grouped aligned kinase domain sequences into hierarchical clusters based on the patterns of functionally conserved residues using the optimal multiple-category Bayesian Partitioning with Pattern Selection (omcBPPS) sampling algorithm (*75*) (see methods for details). In brief, omcBPPS samples the space of sequence partitions and associated sequence patterns to classify aligned protein sequences into multiple hierarchical categories. Sequences that share common co-conserved patterns, also referred to as constraints, are grouped into specific categories or families. The families identified using omcBPPS were then used to build family-specific alignment profiles to refine the classification of bSTK families further. We grouped bSTK sequences that did not fall into any identified families into the "Bac_Unique" category.

We aligned approximately 300,000 bSTK sequences and classified ∼200,000 sequences into 42 newly defined bSTK families (Fig. 1, table S1, data file S2). Taxonomic analysis revealed that many STK families are specific to only one bacterial phylum. Actinobacteria exhibited the most diverse repertoire of STKs, encompassing 13 families and over 100,000 sequences unique to Actinobacterial species. Following this, Proteobacteria comprised 9 families with over 19,000 sequences. Firmicutes included 4 families with 7,000 sequences, Cyanobacteria had three families with 27,000 total sequences, and Myxococcota contained two families with 12,000 sequences. The most prominent bSTK family was KAPD, a gram-positive–specific family that consisted of over 55,000 unique sequences from mainly Actinobacteria and Firmicutes phyla, including one of the most well-studied bSTK proteins, *M. tuberculosis* PknB.

**Fig. 1.**
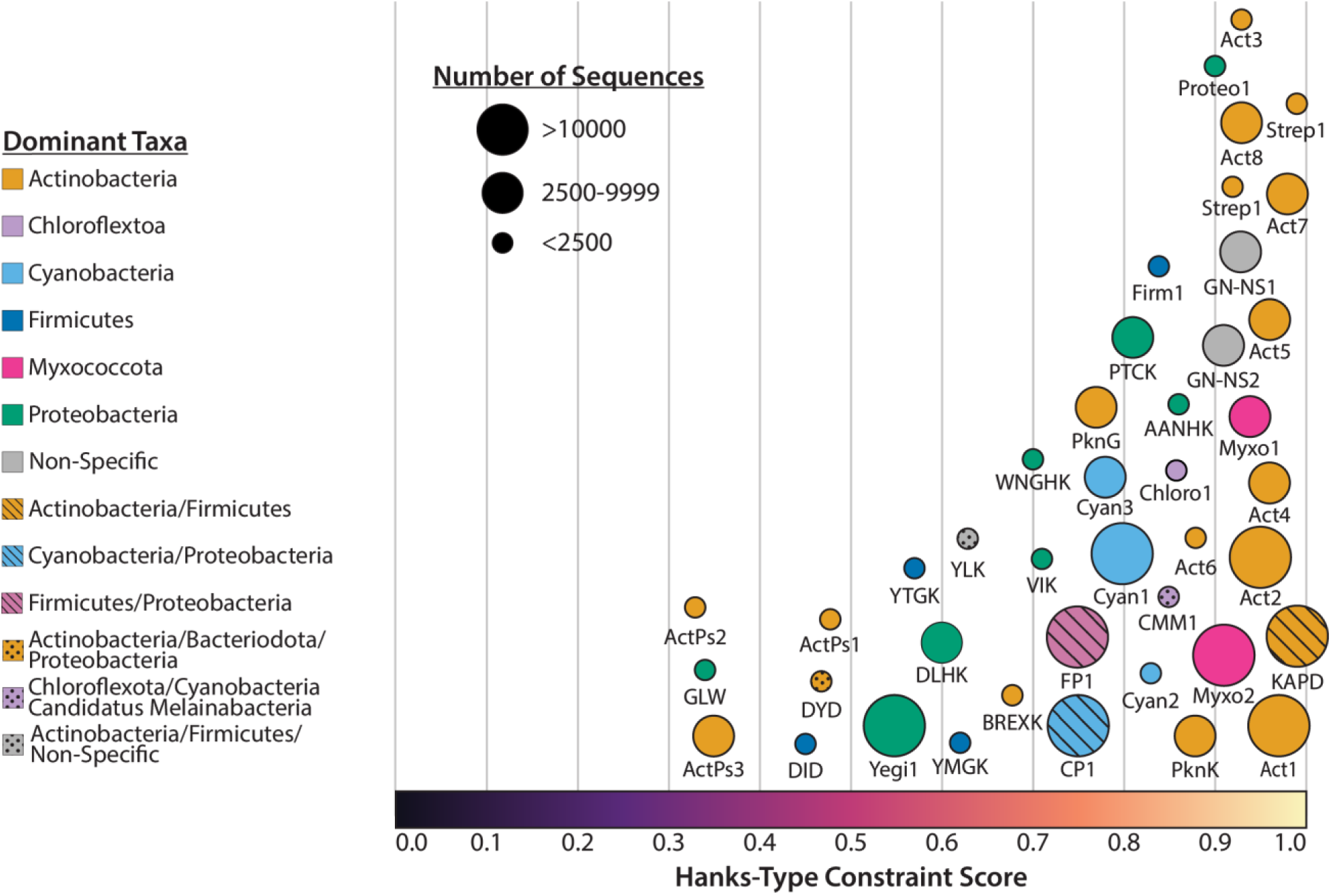
bSTK families identified through constraint-based sequence clustering analysis. Euler diagram of the 42 bSTK families identified using omcBPPS. Families are positioned along the x-axis based on the average Hanks-Type constraint score of the sequences within the family, with 0 representing sequences with no Hanks-Type constraints and 1 representing sequences with all Hanks-Type constraints. The sizes of the families are based on the number of sequences filtered at 98% sequence identity. Acronyms and abbreviations designating the families are defined in the Supplementary Materials (table S2).

### Evolutionarily conserved bacterial pseudokinase families

The Hanks-type kinases are defined by key hallmark function motifs that include the catalytic HRD motif, metal-binding DFG motif, and the salt bridge (K^30^-E^49^, based on the bSTK alignment) connecting the β3 strand and αC-Helix (*4*). Many of the bSTK families scored very highly for the Hanks-type kinase constraints (fig. S1), suggesting that they retain many functionally important Hanks-type kinase motifs and residues. Some bSTK families, however, scored as low as 30% for Hanks-type kinase constraints. The substitution or absence of these constrained residues suggests divergent catalytic or regulatory mechanisms in these families.

Visualization of the sequence patterns (using sequence logos) in these divergent families revealed that seven of the 42 bSTK families displayed variations in one or more catalytic triad residues essential for phosphotransferase activity (Fig. 2A), namely the ATP-binding VAIK-Lys, the catalytic HRD-Asp, and the metal-binding DFG-Asp, and therefore are annotated as pseudokinases (*1*). Substitution of the VAIK-Lys occurs in all seven of the pseudokinase families. 34% of CP1 members, 60% of DYD and all ActPs1, ActPs2, ActPs3, DLHK, and GLW family sequences lack the VAIK-Lys. Substitution of the HRD-Asp occurs in six of the pseudokinase families. 37% of CP1 family members, 97% of the ActPs3, and 99-100% of the ActPs1, ActPs2, DLHK, and GLW family members lack the HRD-Asp. Likewise, multiple bSTK families diverge in the DFG-Asp position. 17% of CP1 family members, 98% of ActPs3, and 100% of ActPs1, ActPs2, and GLW family members lack the DFG-Asp. AlphaFold3 models (*76*) further support the divergence of these families in the active site (Fig. 2B) In addition to the active site, we identified family-specific indels (insertions and deletions) in several pseudokinase families (fig. S2). About 60% of sequences belonging to the ActPs1 family had a deletion of about 20 residues between the F- and H-helices, resulting in the deletion of the G-helix. Similarly, at least 50% of sequences in the ActPs3 family had a 40-residue deletion of the activation loop. The crystal structure of the kinase domain of the MviN protein in *M. tuberculosis*, which falls into the ActPs3 family, further supports this finding by revealing an attenuated activation loop (*77*). ActPs2 sequences, on the other hand, display multiple insertions. The most notable insertions are a small insertion between the end of the activation loop and the F-helix, an insertion of about 30 residues between the β3-strand and the C-Helix, and a ∼200 residue insertion in the activation loop (fig. S2). CP1, DYD, DLHK, or GLW displayed no family-specific insertions.

**Fig. 2.**
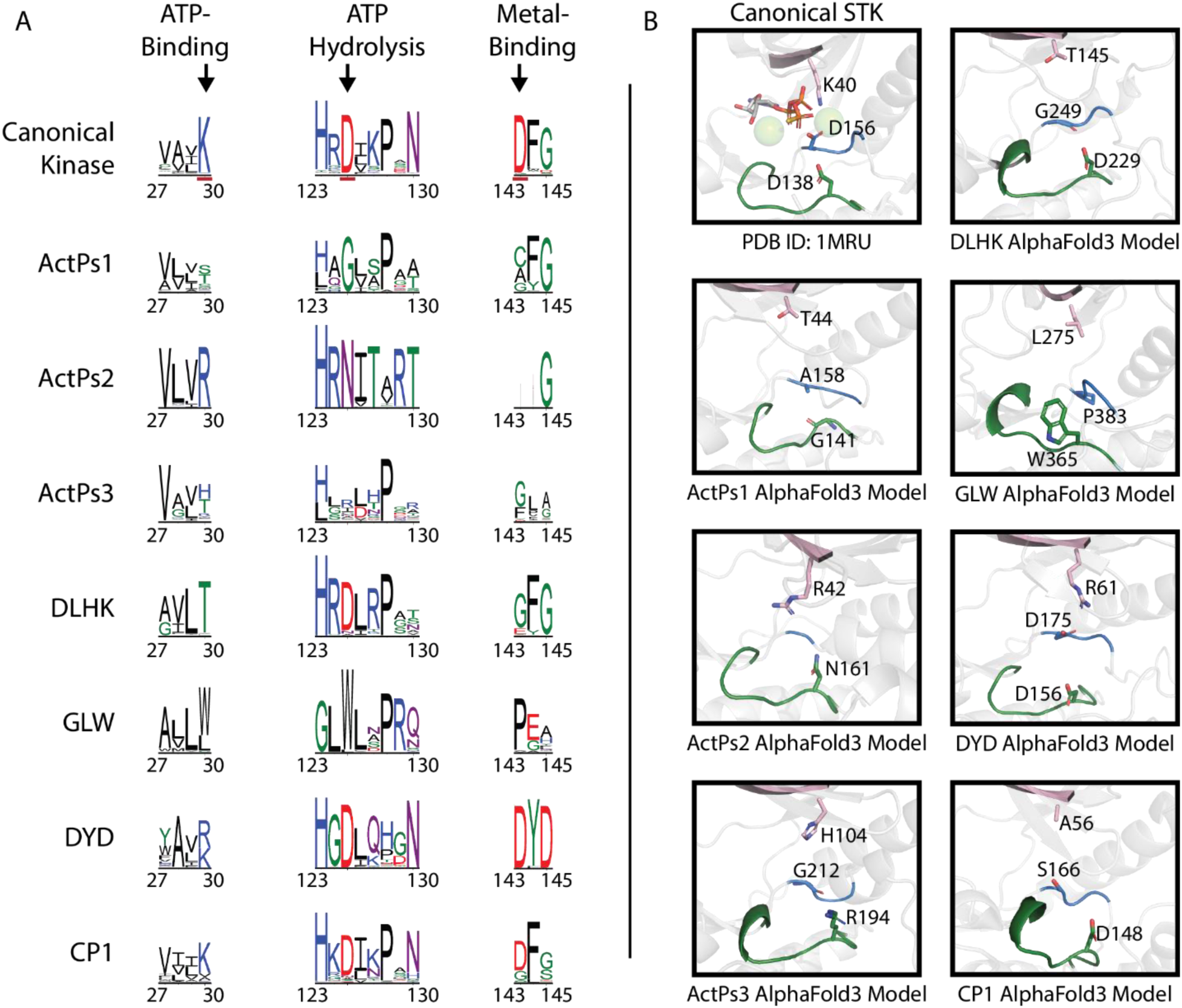
Conserved bacterial pseudokinase families. (**A**) Sequence logos of the catalytic triad motifs in canonical STKs and in each of the bacterial pseudokinase families. (**B**) Comparison between the active site of canonical STKs and the active site of a representative from each bacterial pseudokinase family. The crystal structure of *M. tuberculosis* PknB (PDB: 1MRU) is used as the reference canonical STK. AlphaFold3 models (*76*) were generated for the kinase domain from a representative sequence of each pseudokinase families (NCBI accession numbers: DLHK WP_188258456.1; ActPs1 WP_093894422.1; GLW MBH9021688.1; ActPs2 WP_032763871.1; DYD WP_039364684.1; ActPs3 WP_046703973.1; CP1 WP_248060551.1). The VAIK motif is colored pink, the catalytic loop is colored green, and the DFG-motif is colored blue.

### Phylogenetic reconstruction of bSTK families

After classifying bSTKs into families using constraint-based clustering, we wanted to validate these families and determine evolutionary relationships between them. To accomplish this, we performed maximum-likelihood phylogenetic reconstruction. Following a purge at 100% sequence identity, ∼200,000 protein sequences of bSTKs belonging to the 42 identified families were used in the analysis (data set S2). We found that the phylogenetic analysis corroborated our constraint-based clustering results, with bSTK sequences from the same family clustering together (Fig. 3, data files S3 to 5).

**Fig. 3.**
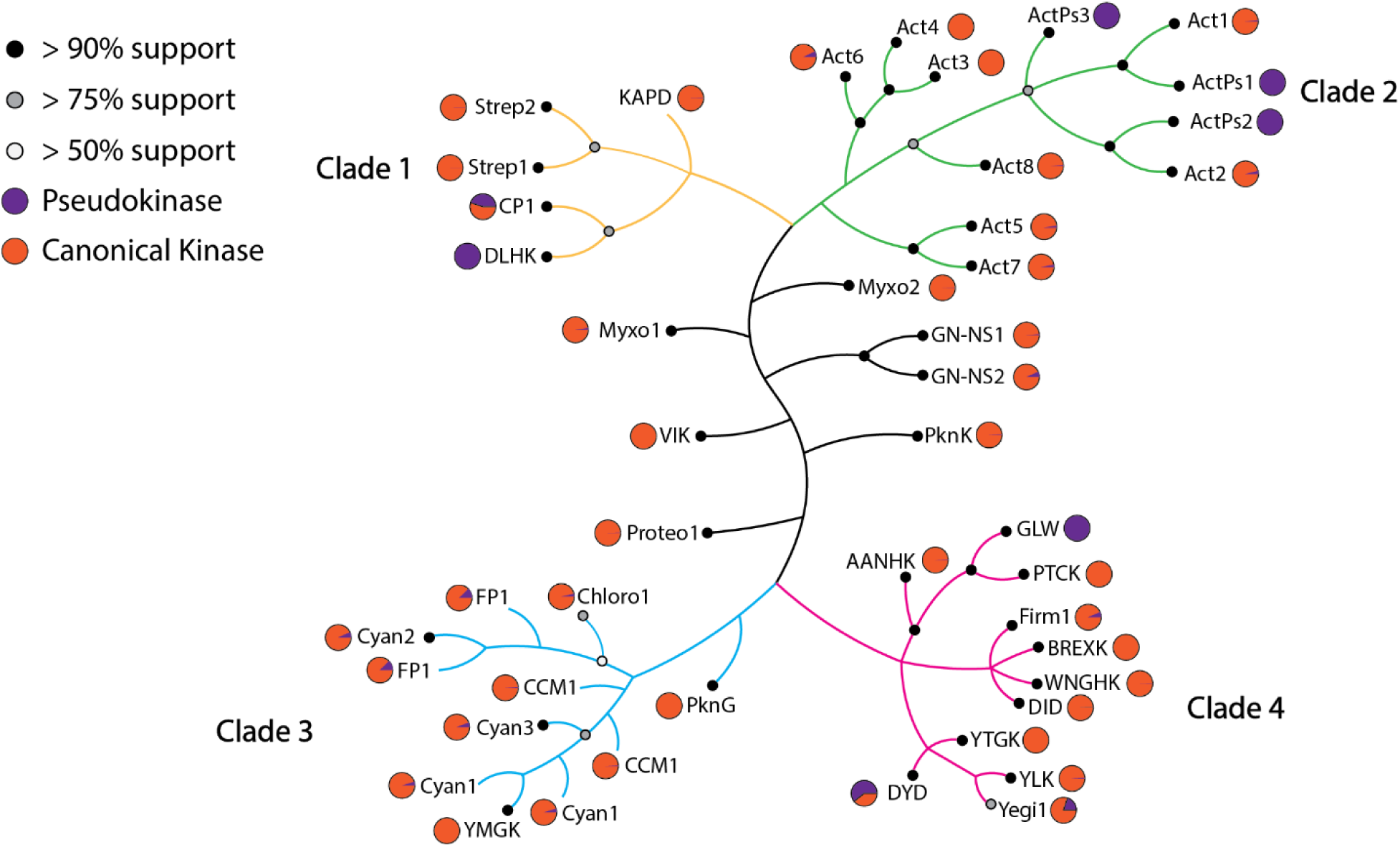
Representation of a maximum-likelihood phylogenetic analysis. The color of internal nodes denotes the branch support of major clades. Lineages belonging to each of the 4 major clades are distinguished by the color of the branches, with Clade 1, 2, 3, and 4 being colored light orange, green, blue, and pink, respectively. Leaf nodes have a pie chart representing the percentage of the canonical kinase (orange) and pseudokinase (purple).

Our phylogenetic analysis reveals that 35 of the 42 bSTK families fall into four major clades Fig. 3, data files S3 to 5): Clade 1, Clade 2, Clade 3, and Clade 4. We found that three of the four clades are taxonomic-specific. 5 families from a diverse array of bacterial phyla make up Clade 1. Clade 2 consists of almost entirely Actinobacteria-specific families, whereas Clade 3 is predominantly Cyanobacteria-specific and includes families from other lineages. Clade 4 is the largest of the four clades and is composed almost entirely of Proteobacteria and Firmicutes-specific families. 7 bSTK families did not fall into any of these four major clades.

Of the 42 families, our phylogenetic analysis revealed that 35 families form monophyletic clades. 6 of the bSTK families are paraphyletic, including Act1, Act2, CCM1, FP1, Cyan1, and PTCK, whereas KAPD is polyphyletic. Act1, Act2, FP1, Cyan1, and PTCK are ancestral to a single other monophyletic STK family (Act1:ActPs1, Act2:ActPs2, FP1:Cyan2, Cyan1:YMGK, and PTCK:GLW). As for CCM1, the family is paraphyletic and split into two groups: (1) Chloroflexota and (2) Cyanobacteria and other phyla. This split is observed across multiple runs regardless of the sequences used, with bootstrap values above 75%.

### Domain and transmembrane analysis reveal family-specific features

We performed domain and transmembrane analysis to obtain functional insights into bSTK families. The most common domain architecture among the bSTK families consisted of a kinase domain with a C-terminal transmembrane helix. We observe this domain organization in 25 bSTK families (Fig. 4). KAPD, GN-NS1, and PTCK families are associated with a second protein domain, in addition to the single STK domain and the C-terminal transmembrane region. These data are consistent with previous domain analyses of bSTK-containing protein sequences that reported bSTKs are linked C-terminally to a transmembrane region (*25*). This contrasts with membrane-associated eukaryotic kinases, which typically have the transmembrane positioned N-terminal to the kinase domain—a domain organization found in only one of the 42 bSTK families, CCM1.

**Fig. 4.**
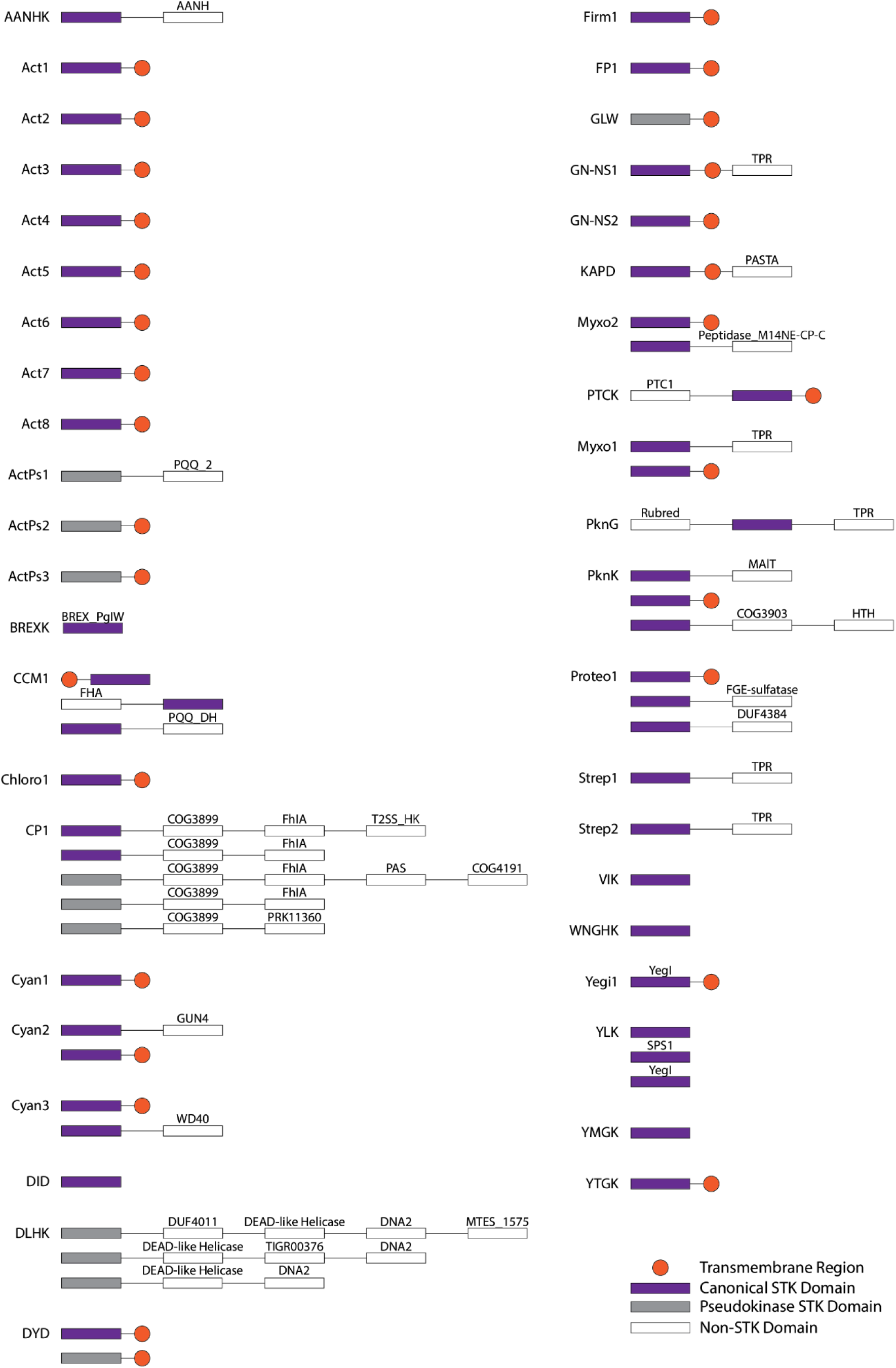
Conserved domain architecture for each of the bSTK families. Canonical STK domains are shown as purple blocks, and transmembrane regions are shown as orange circles. Pseudokinase domains are shown as gray blocks and conserved, non-STK domains are shown as white blocks. Domain architecture is shown for domains present in more than 20% of sequences in the bSTK family. For families containing members with variable domain structures, the different domain organizations are shown. Protein domain acronyms are defined in the Supplementary Materials (table S3).

6 of the bSTK families, CCM1, CP, PknK, Cyan2, Cyan3, Myxo1, Myxo2, Proteo1, and DLHK, are associated with more than one conserved domain architecture. CCM1 family members contain a transmembrane region or an FHA domain N-terminus of the kinase domain. A PQQ DH domain C-terminal of the kinase domain also characterizes a distinct CCM1 subfamily. Likewise, PknK and CP1 family members have multiple functional domains linked to the kinase domain. Cyan2, Myxo1, and Myxo2 families are similar in that they have two possible domain architectures: ones consisting of just a C-terminal transmembrane region and others such as Cyan2, Myxo1, and Myxo2 containing a C-terminal GUN4, TPR, and Peptidase_M14NE-CP-C, respectively. Proteo1 families are associated with three domain organizations: a C-terminal transmembrane region, an FGE-sulfatase, and a DUF4384 domain. DLHK family members generally have four other domains on the same polypeptide chain, all C-terminal to the kinase domain.

5 STK families, Strep1, PknG, Strep2, GN-NS1, and Myxo1, contain conserved TPR C-terminal to the kinase domain. Among all 42 bSTK families, this is the only protein domain that appears alongside multiple families.

### Structural features distinguishing bSTKs from eSTKs

We generated sequence logos from profile-based sequence alignments to investigate the sequence and structural basis for bSTK functional divergence. bSTKs contain multiple unique insertions within the kinase domain, including a 4-residue insertion between the β3-strand and the αC-Helix, a 2-residue insertion in the activation loop, and an insertion of varying length within the F-G-H helical bundle (fig. S3). Overall, bSTK catalytic domain lengths are ∼10 amino acid residues longer than their eSTK counterparts.

We also identified multiple conserved bSTK-specific motifs. These features are highlighted in the *M. tuberculosis* PknB crystal structure (PDB: 612P, Fig. 5A). They include a distinctive methionine in the gatekeeper position (Fig. 5B, fig. S3), a highly conserved methionine in the G-loop (Fig. 5C, fig. S3), a conserved arginine and phenylalanine (RF) motif in the αC-Helix (Fig. 5D, fig. S3), and a poly-proline motif in the loop between the G-helix and the H-helix (Fig. 5E, fig. S3).

**Fig. 5.**
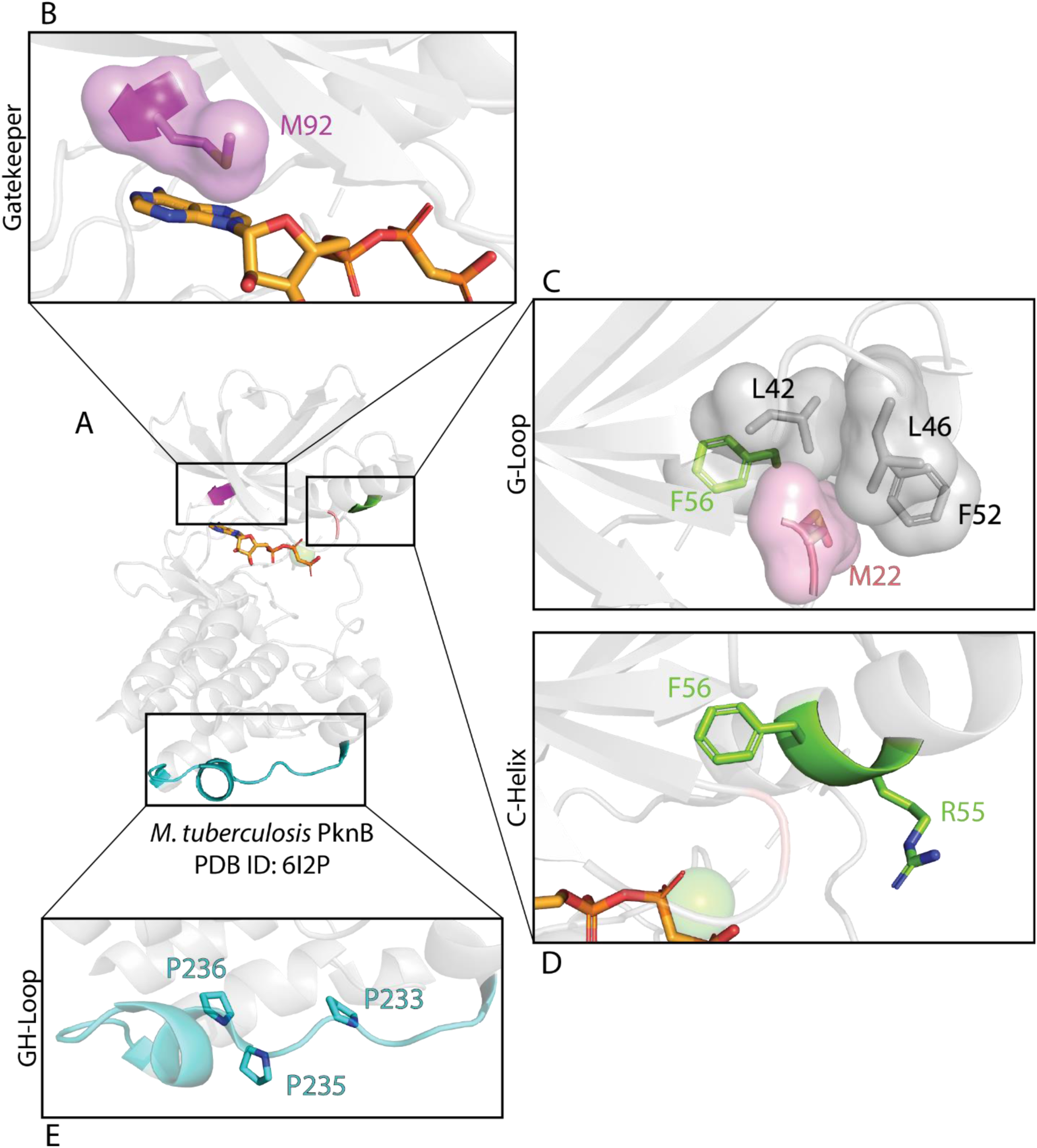
Conserved structural features unique to bSTKs. (**A**) A crystal structure of *M. tuberculosis* PknB bound to ATP and magnesium serves as the reference structure (PDB: 6I2P). (**B**) The gatekeeper-Met is shown in magenta in the reference structure. (**C**) The G-loop-Met is shown in pink in the reference structure, forming a hydrophobic interface with the αC-Helix. (**D**) The RF motif located within the αC-Helix is colored green. (**E**) The prolines of the poly-proline motif of the GH-loop are shown in cyan.

One of the most distinguishing bSTK features is an RF motif in the αC-Helix (Fig. 5D). The RF-Phe mediates hydrophobic packing interactions with residues in the N-lobe, particularly a Met in the G-loop. The arginine (Arg^45^ based on the bSTK alignment) is present in ∼50% of bSTKs (Fig. 6A) and highly conserved in 22 bSTK families (fig. S4). In contrast to the RF-Phe, the RF-Arg is quite dynamic and adopts multiple distinct conformations in available crystal structures and AlphaFold3 models (Fig. 6B). We group these conformations into five distinct states. In State 1, the C-Helix-Arg interacts with the phosphate groups of phosphorylated threonines in the activation loop (Fig. 6C), whereas in State 2 it forms a cation-pi interaction with a phenylalanine of the C-Helix (Fig. 6D). In State 3, the C-Helix-Arg forms a salt bridge with the highly conserved C-Helix-Glu (Fig. 6E), whereas in State 4 it hydrogen bonds to the backbone carbonyl of the DFG-Gly in the activation loop (Fig. 6F). In the substrate-bound structure of PknB, the C-Helix-Arg forms a salt bridge with a glutamate in the Glycogen accumulation regulator A (GarA) substrate (Fig. 6G).

**Fig. 6.**
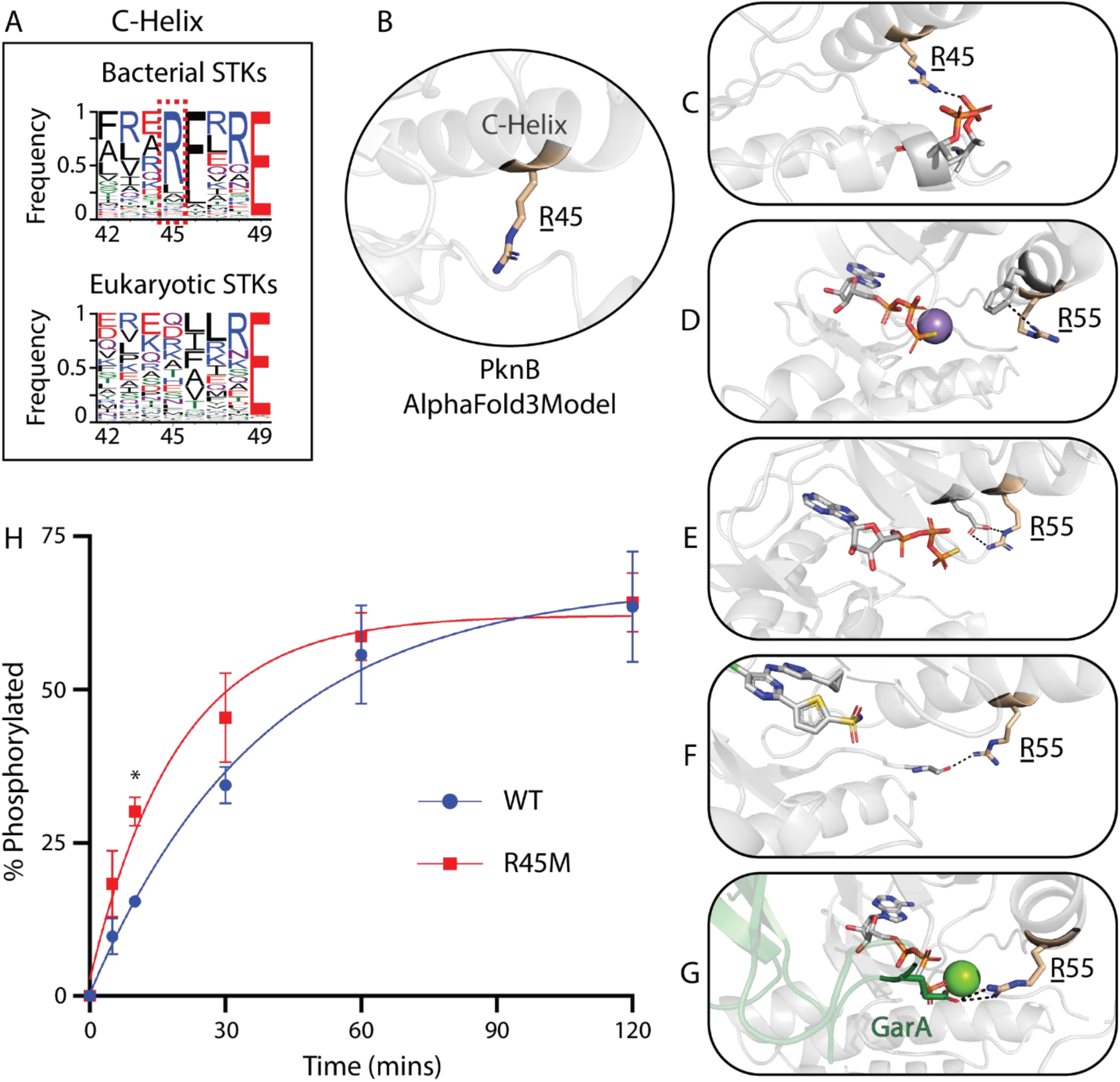
Conservation and dynamic nature of the C-Helix-Arg in bSTKs. (**A**) Sequence probability logo of the canonical bSTK and eSTK C-helices. The conserved, bacterial-specific C-Helix-Arg is annotated with a dotted red box. (**B**) AlphaFold3 model of the C-Helix-Arg, shown in light brown, without interactions. (**C**) AlphaFold3 model of C-Helix-Arg interaction with phosphorylated threonine residues in the activation loop. (**D**) Crystal structure (PDB: 3F69 Chain B) of the C-Helix-Arg forming a cation-pi interaction with a phenylalanine within the C-Helix. (**E**) Crystal structure (PDB: 3ORT) of the C-Helix-Arg forming a salt-bridge with the C-Helix-Glu. (**F**) Crystal structure (PDB: 6B2P) of the C-Helix-Arg forming a hydrogen bond with the backbone of a residue within the activation loop. (**G**) Crystal structure (PDB: 6I2P) of the C-Helix-Arg (Chain B) forming a salt-bridge with a co-crystalized substrate, GarA, shown in dark green. (**H**) Rate of in vitro autophosphorylation of His-tagged WT and R45M PknB at different time points (0, 10, 30, 120 minutes). The percent phosphorylation at each time point was determined based on the phosphorylated and unphosphorylated protein band densities quantified using a LICOR Odyssey M imager (fig. S5). Autophosphorylation assays were performed in triplicate and significance was calculated using an unpaired t-test for each time point (*, *P*<0.05).

To evaluate if the C-Helix-Arg plays a role in modulating kinase activity, we mutated the *M. tuberculosis* PknB C-Helix-Arg^55^ (referred to as Arg^45^ for consistency) to a methionine (R45M), a variant observed on other bSTKs (fig. S4) and measured the impact of R45M mutation on kinase autophosphorylation activity. Autophosphorylation activity is seen almost immediately in both PknB variants following the addition of Mg-ATP (Fig. 6H and fig. S5). However, after 10 minutes the R45M autophosphorylation rate was significantly faster than WT, with almost double the percent phosphorylation. The R45M autophosphorylation rate remained faster than WT for the first 30 minutes of the assay and only after 60 minutes did both the WT and R45M variants display comparable amounts of autophosphorylation. These results suggest a role for R45 in kinase regulation.

### Effects of constraint mutations on PknB substrate specificity motif

To demonstrate that the bSTK constraints have biological relevance in addition to being hallmark features of their respective families, we mutated the *M. tuberculosis* PknB C-Helix-Arg^45^ and a KAPD-specific constraint within the D-helix, Arg^101^ (referred to as Arg^87^ for consistency), to a methionine (R45M) and alanine (R87A), respectively. These amino acid variants are found in other bSTKs and were selected to preserve the activity and stability of the kinase. We then used a Positional Scanning Peptide Array (PSPA) analysis to generate a substrate specificity consensus motif for the Wild-Type PknB and the two mutants (data files S6 to S8). The wild-type (WT) PknB strongly prefers negatively charged residues and has a low tolerance for hydrophobic residues N-terminus of the phosphoacceptor site (Fig. 7A, fig. S6). On the C-terminus of the phosphoacceptor site, the WT PknB generally prefers charged or polar residues, particularly at the +1 position, where the most highly preferred residue is glutamine. In contrast, the R45M mutation substantially altered peptide substrate preference. Although amino-acid preferences N-terminus of the phosphoaccetor site are comparable to WT (Fig. 7B, fig. S7), consensus motifs in the C-terminus are substantially different, with preference for hydrophobic residues at the +2, +3 and +4 positions and some preference for a polar or charged amino acid at the +1 position. In the case of the R87A mutant, the substrate specificity consensus motif is drastically different from WT on both termini of the phosphoacceptor (Fig. 7C, fig. S8). On the N-terminal side of the phosphorylation site, the amino-acid preference changes from acidic to basic or polar amino acids and demonstrates an increased tolerance for hydrophobic residues. Overall preference for specific residues N-terminus of the phosphorylation site is lower than WT. On the C-terminus of the phosphorylation site, however, R87A shows a strong preference for hydrophobic residues, particularly at the +2 and +3 positions, and a negative selection for acidic/charged residues and prolines.

**Fig. 7.**
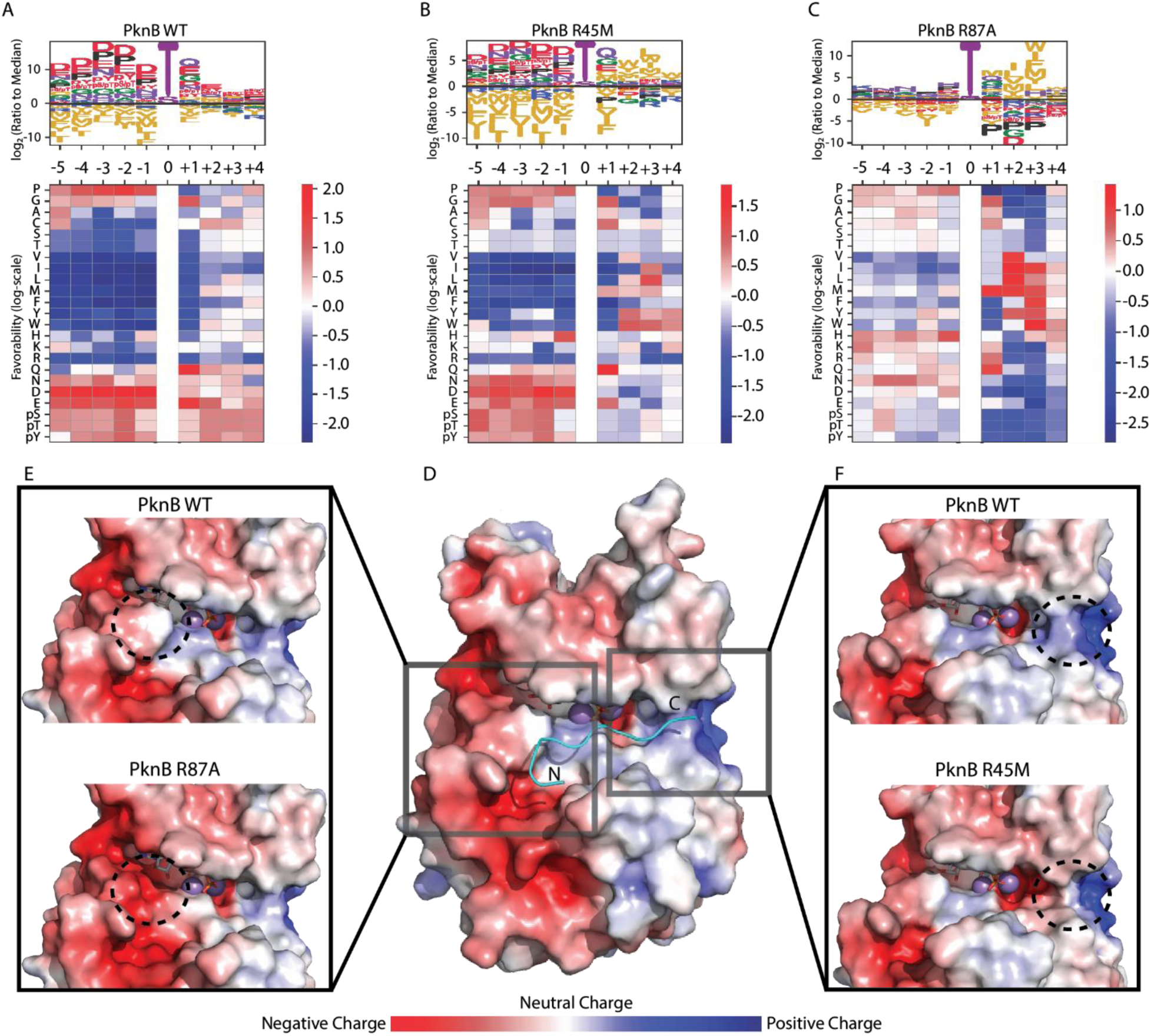
Substrate specificity consensus motif of *M. tuberculosis* PknB mutants and their modifications to surface electrostatics. (**A**) PknB WT substrate specificity consensus motif and heatmap. The effect that an amino acid at a given position within the peptide has on substrate phosphorylation relative to other amino acids at the same position is noted with larger characters in the sequence logo indicating a more substantial effect on phosphorylation. Amino acids above the x-axis indicate a positive effect on the favorability of substrate phosphorylation relative to other amino acids, and amino acids below the x-axis indicate a negative effect on favorability. The heatmap provides the log-scale favorability values for each amino acid at each position within the peptide, with indicating positive favorability values and blue indicating negative favorability values. (**B and C**) substrate specificity consensus motifs and heatmaps for PknB R45M (B) and PknB R87A (C). (**D**) Surface electrostatic visualization of PknB WT bound to a modeled peptide (shown in cyan). N- and C-termini of the peptide are indicated with N and C, respectively. Red indicates a negative charge, blue indicates a positive charge, and gray indicates a neutral charge. (**E and F**) Comparison of the PknB WT surface electrostatics with those of PknB R45M (E) and PknB R87A (F). The dashed circles indicate the arginine side chain in the WT protein and its absence in the mutant proteins to highlight its effect on the local surface electrostatics.

To obtain structural insights into the altered substrate specificity of the R45M and R87A mutants, we modeled a peptide into the substrate binding region of PknB and visualized the surface electrostatics (Fig. 7D). The R87A mutation creates a negatively charged surface patch surrounding the D-helix relative to WT PknB and a deeper groove for accommodating residues in the N-terminal region of the substrate peptide (Fig. 7E). In contrast, the R45M mutant creates a more neutral charge in a large pocket near the junction of the C-Helix and the activation loop, near where the C-terminus of the peptide binds (Fig. 7F). Together these structural changes provide a plausible explanation for the observed differences in the peptide substrate specificity profiles displayed by mutant and WT PknB.

## Discussion

In this study, we report a comprehensive classification of bSTKs into 42 distinct families based on a comprehensive informatics analysis of over 300,000 bacterial kinase sequences. We demonstrate that bSTKs are a diverse class of enzymes with substantial variations in both primary sequence and 3D structures. The constraint patterns represent the defining residues for each bSTK family and serve as a quantitative metric for annotating sequences into one of the families. These constraints can be utilized to characterize family-specific regulatory and mechanistic traits at the molecular level. We demonstrated this in PknB, in which the mutation of KAPD-specific arginine (R^87^ in reference to the bSTK alignment) in the D-helix results in a considerably different substrate specificity consensus motif compared to the WT PknB.

Actinobacteria have the most diverse array of bSTK families. This phylum is one of the largest, most common, and most diverse bacterial phyla, with members of this taxonomic group represented in terrestrial and aquatic ecosystems (*78*). Previous studies have reported that in Actinobacteria, protein kinases are the predominant type of membrane-associated signaling component (*79*). The diversity of Actinobacterial STKs identified in our study further supports their proposed roles in diverse bacterial functions (*78*, *80*).

Although the functions of most bSTKs are yet to be determined, our analysis of their domain architectures suggest important functional roles. For example, the KAPD family of STKs carry a conserved penicillin-binding and serine-threonine kinase–associated (PASTA) domain, C-terminal to the kinase domain and a transmembrane region. This architecture is well-documented in bSTK, with multiple PASTA domains in the same polypeptide chain (*81*). In *Streptococcus pneumoniae* and *M. tuberculosis,* the PASTA domains serve as extracellular sensors by binding peptidoglycans and subsequently activating their associated STK, which phosphorylates its substrate and begins a signaling cascade to regulate cell division or peptidoglycan synthesis (*30*, *82–84*). Given the widespread presence of membrane-associated bSTK families (28 of 42 bSTK families), many of the STK families may be involved in communicating extracellular signals to intracellular response in a manner analogous to receptor tyrosine kinases. Another domain of note is the TPR domain, which is present in multiple divergent bSTK families mapping to different clades in the phylogenetic tree. TPR domains function as protein scaffolding or interaction modules in many biological functions, including protein import, gene regulation, and mitosis (*85*). TPRs are also implicated in host cell adhesion, virulence factor transportation, and the survival of pathogenic organisms within a host (*86–89*). The crystal structure of PknG, a secreted STK from *M. tuberculosis*, suggests a role for TPR domain in protein dimerization (*90*). We predict that the TPR domain performs an analogous protein interaction role in the GN-NS1, Myxo1, Strep1 and Strep2 families identified in our study.

Our study also maps and extends the pseudokinase complement of bSTKs. Conserved eukaryotic pseudokinase families perform important noncatalytic functions such as protein scaffolding, competitive inhibitors, molecular switches, and allosteric activators (*25*, *91*, *92*). And although their presence in bacteria is not as ubiquitous, the presence of pseudokinase STK families within bacteria and their unique constraint patterns not seen in eukaryotic kinases hint towards conserved, non-catalytic activity of these families, potentially performing bacterial-specific roles. In eukaryotes, some pseudokinase families also retain the ability to bind ATP or non-canonical small molecules (*93*) despite lacking the VAIK-Lys, such as the WNK family (*94*), ULK4 (*18*), or KSR1 (*95*, *96*). Given the conserved substitution of the VAIK-Lys to an arginine, ActPs2 sequences may have evolved non-canonical ligand binding mechanisms (Fig. 2). In DLHK family members, the presence of the HRD-Asp, which is required for ATP hydrolysis, suggests that members of this family may retain the ability to hydrolyze ATP; however, the substitution of both the metal-binding DFG-Asp and HRDxxxxN-Asn would also suggest non-canonical nucleotide binding

bSTKs display unique sequence and structural variations in the catalytic domain that provide clues for functional inference and drug development efforts. One of the most distinctive features distinguishing bSTKs from eSTKs is the RF motif in the regulatory C-Helix. The RF motif is rarely observed in eSTKs with the exception of some apicomplexan STKs, which conserve an arginine structurally analogous to the RF-Arg in bSTKs (*97*). In the apicomplexan kinase CpCDPK2 (*Cryptosporidium parvum* Calcium Dependent Protein Kinase 2), the C-Helix-Arg coordinates with the phosphorylated residue in the activation loop similar to the phosphorylated PknB model proposed in our study (Fig. 6C). Furthermore, the multiple conformational states adopted by the RF-motif arginine in different crystal structures of PknB suggests an auto-regulatory role for this motif in bSTKs. Indeed, mutation of the RF-Arg to a methionine alters the rate of autophosphorylation, presumably by relieving the autoinhibitory interactions between the RF-Arg and the C-Helix-Glu that prevents the kinase from adopting an active conformation (Fig. 6E). In this regard, the bSTK-specific insert between the β3 strand and the C-Helix (fig. S3) may be relevant because this region is also known to modulate C-Helix movement in PknB and in eSTKs (*98*).

Another striking feature of bSTKs is the divergence in the ATP binding gatekeeper residues and substrate binding GHI sub-domain. Most eSTKs have a hydrophobic residue or a threonine at the gatekeeper position (PKA Met^121^), though it varies based on the family. Across bSTKs, however, there is a substantially lower amount of diversity at the gatekeeper position, with methionine being the predominant residues at the gatekeeper in bSTKs. This information can be leveraged to inform drug repurposing efforts aimed at regulating bSTK activity, as the properties of the gatekeeper residue dictate nucleotide binding and Type-1 kinase inhibitor selectivity (*99*). A methionine-selective Type-1 kinase inhibitor would serve as a conceptual starting point for bSTK-specific inhibitor design. Likewise, we identified a left-handed poly-proline type II (PPII) helix motif within the GHI-subdomain of some bSTKs, a feature that is not seen in eSTKs. PPI helices have been shown to serve as protein docking sites (*100*) and the GHI domain is known to play a role in protein-protein interactions and substrate binding for both bSTKs and eSTKs (*98*, *101*). With this information, one can identify potential bSTK substrates and binding partners, further characterizing the functions of bSTKs in cellular signaling pathogenicity and informing the development of bSTK targeting drugs.

A structure of PknB cocrystalized with a substrate, GarA, also reveals the C-Helix-Arg plays a role in substrate specificity, which is further corroborated by our Positional Scanning Peptide Array analysis. An arginine to methionine mutation results in PknB preferring a substrate with hydrophobic residues C-terminal to the phosphorylation site. Our analysis also revealed that bSTKs bind peptide substrates in a canonical orientation and that the KAPD-specific constraint, Arg^87^, plays a role in substrate specificity, as previously suggested (*102*). Arg^87^, found within the D-Helix, may only require a single acidic residue N-terminal of the phosphorylation site, with the arginine side chain’s flexibility enabling it to interact with any residue at the five N-terminal positions. This would explain why there is such high tolerance for non-acidic residues at those positions within the peptide, and why a single substitution resulted in the loss of specificity towards acidic residues at all five N-terminal positions. Unlike the R45M mutation, however, the R87A not only affects local specificity on the N-terminal side of the phosphoacceptor site, but also the specificity of more distal residues C-terminus to the phosphoacceptor site. This degree of change in the substrate specificity consensus motif is substantial, especially for a single amino acid substitution. This includes an increase in the favorability for methionine at the +1 position over glutamine, which previously was suggested as a crucial residue for PknB substrate phosphorylation (*49*). The exact mechanism for how Arg^87^ affects the specificity for distant residues of the peptide remains unclear, however, it further emphasizes the importance of these bSTK family constraints in the structure-function and specificity of bSTKs.

bSTKs have been shown to play important regulatory roles in bacterial pathogens, making them attractive targets for anti-microbial therapeutics (*30*, *57*, *103*, *104*). Several studies have explored this avenue in an effort to combat antimicrobial drug resistance (*105–109*), however, there are currently no bSTK inhibitors approved for antimicrobial use (*110*) due to issues such as toxicity. Many pipelines and assays are already in place to develop therapeutics that target human STKs (*111–114*). Tailoring these to bSTKs would provide a robust framework for drug development using existing resources, and the constraints that define each of the bSTK families would provide clues for repurposing existing inhibitors to target bSTKs with a greater degree of specificity. The structure and regulatory features we have presented in this paper that are shared among bSTKs, such as bSTK-specific divergence in the gatekeep residues and ATP binding P-loop, could also be used to inform the development of broad spectrum bSTK targeting antimicrobial drugs.

## Materials and Methods

### Detection of bSTK sequences

In order to align and annotate bSTK sequences from protein databases, 43 experimentally validated bSTK sequences (data file S1) were gathered and aligned using MAFFT v7.453 to create a Multiply-Aligned Profiles for Global Alignment of Protein Sequences (MAPGAPS) profile. Sequences from closely related kinase families identified in Kannan, 2007 (*24*) were also aligned and included in the MAPGAPS profile to prevent the misannotation of non-protein kinases as bSTKs. MAPGAPS version 2.1 was then used to annotate bSTKs by aligning protein sequences from the UniRef protein sequence database (obtained February 6, 2020) to the bSTK MAPGAPS profile, from which 45,359 bSTK sequences were annotated. Following an initial run of omcBPPS, protein sequences that fell into STK families identified through omcBPPS were used to build a hierarchical MAPGAPs profile in an effort to further refine protein sequence alignment and STK identification. The NCBI Non-Redundant protein sequence database (obtained June 14, 2022) was also aligned to later versions of the bSTK profile with a total of 500,935 bSTK sequences identified, 294,534 of which were annotated into one of the 42 bSTK families we identified.

### Bayesian statistical analyses and pattern-based classification of bSTK sequences

Constraint-based sequence clustering was performed with the Optimal Multi-Category Bayesian Partitioning with Pattern Selection (omcBPPS) scheme and used to group STKs together into families of evolutionarily related proteins. All omcBPPS runs were run in triplicate. For the initial run, 45,359 bSTK sequences that were initially identified using the MAPGAPS algorithm were used as the input for omcBPPS. Sequences that contained constraint patterns identified from the omcBPPS run were used to build a MAPGAPS profile for their corresponding family. We aligned sequences from the aforementioned protein sequence databases using the MAPGAPS algorithm. The sequences that were annotated by MAPGAPS to belong to each bSTK constraint-cluster (family) were then purged at 95% sequence identity (42,543 sequences) and used as input for omcBPPS to further breakdown each constraint-cluster into unique families or subfamilies. bSTK sequences that did not belong to any of the constraint clusters were also purged at 95% sequence identity and used as an input for omcBPPS to sort them into newly classified bSTK families. This process was repeated multiple times, using HelperBunny to identify whether a constraint pattern matched a previously identified pattern or represented a previously undescribed pattern.

We used a quantitative means of evaluating how well any given sequence fit into each of the clusters defined by our model, as previously performed in Yeung et al, 2021 (*65*). Each bSTK cluster was associated with a list of constraints that dictated which amino acid(s) were likely to be found at a given alignment position. Furthermore, each constraint was associated with a log-likelihood score which described how specific a constraint was to its respective cluster. To score a sequence against a cluster, we added the log-likelihoods of all the constraints which were true for the query sequence. In order to make this score comparable across clusters, we divided this number by the total log-likelihood of all the cluster’s constraints. This resulted in a range from 0 to 1. For example, a sequence which followed all of a given cluster’s constraints would have received a maximum score of 1 for that cluster, whereas a sequence which did not satisfy any of a given cluster’s constraints would have received a minimum score of 0 for that cluster. For the purposes of discrete classification, we defined a cut-off score for classifying a sequence into a cluster. Based on the distribution of scores from all possible sequence-cluster pairs across multiple test data sets, we defined the optimal cut-off at the global minimum of 0.7 (fig. S9).

### Comparative Sequence Analysis

Comparative sequence analyses between bSTK families and EPKs were performed in Python3 using custom scripts (provided in our computational notebooks). Weblogos were generated using WebLogo 3.7.12, with the colors of the amino acids corresponding to their biochemical properties.

### Modeling protein structures

Protein models for STKs that did not have protein structures available in the Protein Data Bank were predicted using AlphaFold version 3 (*76*). Structures were visualized in PyMOL version 2.3.0 (*115*). Constraints associated with the bSTK families were mapped to AlphaFold models or crystal structures using a custom Python3 script (provided in our computational notebooks).

Substrate bound *M. tuberculosis* PknB structure was generated through homology modeling using SWISS-MODEL (*116*). PkA bound to PKI was used as the homology model template (PDB: 2GFC) and the canonical *M. tuberculosis* PknB kinase domain (UniProt: P9WI81 residues 11-273) was used as the query sequence.

### Phylogenetic reconstruction

Phylogenetic reconstruction of aligned bSTK domains was performed to model STK evolution in bacteria. The phylogenies of larger datasets were inferred through maximum-likelihood using FastTree2.1 (*117*). All bSTK sequences from the NCBI sequence database belonging to the 42 identified families were first purged at 100% sequence identity using CD-HIT (*118*) and outlier sequences were removed (201,421 sequences total). FastTree2 was run using the LG model and Gamma20 likelihood was optimized following CAT (categories) approximation. Local support values were calculated using the Shimodaira-Hasegawa test (*117*).

Phylogenetic analyses were performed on smaller datasets to validate the FastTree results. Phylogenies of datasets containing representatives from each of the bSTK families were inferred using IQ-TREE version 1.6.12 (*119*) with ModelFinder (*120*). bSTK families were first purged at 90% sequence identity and 15 representative sequences were randomly selected from each family. 5 EPK sequences were also included in the dataset to serve as an outgroup. The substitution model LG (*121*) was used with empirical amino acid frequencies and a FreeRate model with 10 categories (*122*, *123*). Branch support values were generated using ultrafast bootstrap with 1000 resamples (*124*, *125*). Results indicated that sequences from the same clusters consistently formed paraphyletic groups or monophyletic clades with high bootstrap support.

### Domain and transmembrane analysis

Transmembrane regions were predicted by running full length protein sequences through TMHMM-2.0 (*126*). Additional domains for each bacterial protein sequence were identified using the bwrpsb api on NCBI with superfamily hits only against the CDD database (*127*) with an e-value threshold of 0.01 and with a maximum number of 500 hits. Kinase domains were annotated with our manually curated protein kinase profiles and any domains overlapping the kinase domain were removed.

### Protein expression and purification

The PknB construct (11-288aa) was codon optimized for expression in *Escherichia coli* using the GeneSmart prediction software (Genscript). DNA sequence was cloned into a pET-28a(+) expression vector at cloning sites EcoRI/NotI.

The plasmid containing N-terminal His-tagged PknB was expressed in E. coli BL21 (DE3) cells. Bacteria were cultured in LB media supplemented with 60 µg/mL kanamycin at 37°C. Upon reaching an optical density (OD) of 0.6, E. coli was induced with 1 mM IPTG (Isopropyl β-D-1-thiogalactopyranoside) for 16 hours at 15°C. Subsequently, the bacterial culture was centrifuged at 5000 rpm at 4°C for 15 minutes. The resulting bacterial pellet was then resuspended in a buffer consisting of 50 mM HEPES, 200 mM NaCl, 10% glycerol, and 1 mM PMSF (phenylmethylsulfonyl fluoride) at pH 7. The lysate was clarified through centrifugation at 15,000 rpm at 4°C for 1 hour. The resulting supernatant, containing solubilized PknB, was collected and vacuum filtered through a 0.45 µm filter unit (Fisher Scientific). The bacterial lysate was loaded onto a Cytiva HiTrap TALON crude 1 mL column and eluted across a 0-500 mM imidazole gradient in 1 mL fractions. Protein purity was assessed using SDS-PAGE with Coomassie stain. Fractions were collected and dialyzed against a buffer consisting of 50 mM HEPES, 200 mM NaCl, and 10% glycerol at pH 7 four times over 20 hours. Quantification of PknB was performed using the Bradford Assay (Bio-Rad), revealing a concentration of 0.85 mg/mL. Protein stability was assessed through Differential Scanning Fluorimetry (Applied Biosystems StepOnePlus Real-Time PCR System) for 2 hours with a step increase from 25°C to 95°C. Protein function was assessed using an autophosphorylation assay. The protein was then aliquoted, snap-frozen, and stored at −80°C.

R81M (equivalent to R45M within the bSTK alignment and R55M within the *M. tuberculosis* PknB P9WI81 UniProt sequence entry) and R127A (equivalent to R87A within the bSTK alignment and R101A within the *M. tuberculosis* PknB P9WI81 UniProt sequence entry) mutations in the PknB kinase domain were introduced using the New England Biolabs Q5 site-directed mutagenesis kit following the manufacturer’s recommendations. Primers were designed using the online NEBaseChanger tool and were generated by Eurofins.

### Autophosphorylation Assay

WT PknB and R45M PknB (both 0.8 mg/mL) were dephosphorylated by incubating with New England Biolabs Lambda Phosphatase, 10mM MnCl_2_, and 10x NEB buffer at room temperature for one hour. To separate the His-tagged kinase from the untagged phosphatase, Co-NTA agarose beads were added to the reaction tube and incubated at 4°C for 1 hour. The beads were washed three times with buffer (50mM HEPES, 200mM NaCl, and 10%, pH 7) and the bound proteins were eluted in wash buffer containing 300mM imidazole. Eluted protein was dialyzed in wash buffer to remove imidazole. Quantification of WT PknB and R45M PknB was performed using the Bradford Assay (Bio-Rad) and the final concentration for both variants was 0.5 mg/mL.

For the autophosphorylation assay, 2x kinase buffer (50mM HEPES, 200mM NaCl, 10% glycerol buffer, 20mM MgCl_2_, pH 7) and 5mM ATP were added to a microcentrifuge tube containing either PknB WT or PknB R4M. Tubes were put in a 30°C water bath and samples were taken from the reaction tube at 0, 5, 10, 30, 60 and 120 minutes and the reaction was immediately quenched with 4x SDS reducing sample buffer and 5 minutes on a 95°C heat block. Samples were loaded on a 10% SDS-PAGE, run at 151V for 1 hour and stained with Coomassie blue. The density of the bands of phosphorylated and dephosphorylated protein were quantified using a LICOR Odyssey M instrument.

### Profiling kinase substrate specificity

Recombinant *M. tuberculosis* PknB preferred substrate was identified using a Positional Scanning Peptide Array analysis, performed as outlined in Johnson et al., 2023 (*14*). PknB was added to a 384-well plate that contained a 50uM solution of peptide substrate library mixtures (Anaspec, AS-62017-1 and AS-62335). Assay conditions were as follows: 50 mM HEPES, 200 mM NaCl, and 10% glycerol at pH 7.3. The reaction was initiated with 100 uM of ATP (50 μCi ml−1 γ-32P-ATP, Perkin-Elmer), followed by a 90-minute incubation. After the reaction, solutions were spotted onto streptavidin-conjugated membranes (Promega, V2861), where the biotinylated C-terminal ends of the peptides were tightly associated. The membranes were rinsed, and peptide phosphorylation amounts were imaged using a Typhoon FLA 7000 phosphorimager (GE). Densitometry matrices were generated by quantifying the raw data (GEL file) using ImageQuant (GE).

## Supplementary Materials

Figs. S1–S9.

Tables S1–S3.

Data files S1–S8.

## Acknowledgements

Thank you to members of Dr. Natarajan Kannan’s laboratory, Dr. Patrick Eyers, Dr. Dominic Byrne, and Dr. Benjamin Turk for helpful discussions and valuable feedback.

## Author contributions

Conceptualization: BMO, NK

Data curation: BMO, WY

Formal analysis: BMO, WY, JLJ, TMY

Funding acquisition: NK

Investigation: BMO, JDL, SK, JLJ

Methodology: BMO, WY, JDL, SK, NK

Project administration: NK

Resources: NK LCC

Supervision: NK, SK

Validation: BMO, JDL, JLJ

Visualization: BMO, TMY

Writing – original draft: BMO, JDL, NK

Writing – review & editing: BMO WY, JDL, JLJ, NK

## Data and materials availability

Jupyter notebooks and full protein sequence datasets are available at the following github repository: https://github.com/boboyle/Atlas-of-Bacterial-Serine-Threonine-Kinome.git. All other data required to evaluate the conclusions in the paper are present in the paper or the Supplementary Materials.

